# Older Adults Show Preserved Neural Signatures of Audiovisual Integration but Altered Links to Speech Comprehension

**DOI:** 10.64898/2026.10.26.761765

**Authors:** Charlie Reynolds, Ole Jensen, Katrien Segaert, Hyojin Park

## Abstract

Older adults often experience difficulty understanding speech in noisy environments despite relatively preserved hearing. Although visual speech cues can improve speech perception, it remains unclear whether the neural mechanisms supporting audiovisual integration are preserved in healthy ageing. We addressed this question using magnetoencephalography (MEG) combined with Rapid Invisible Frequency Tagging (RIFT) during challenging audiovisual speech task. Forty younger and forty older adults attended to one of two competing speech streams while the speaker’s lip movements were frequency tagged, with the visual modulation driven by either the attended or unattended speech envelope. Task-Relevant amplitude modulation of frequency tagging in visual speech improved speech comprehension in younger adults as previously reported for this cohort; this accuracy benefit was absent in older adults. Auditory and visual tagging responses were modulated by task relevance similarly across age groups, indicating largely preserved modality-specific sensory processing in older adults. Intermodulation responses, reflecting nonlinear interactions between auditory and visual speech signals, also remained sensitive to task relevance and did not differ in their condition-dependent modulation between age groups. Greater Task-Relevant versus Task-Irrelevant differences in intermodulation coherence were also associated with greater comprehension benefits in younger adults; no significant association was found in older adults, and this relationship differed significantly between age groups. These findings suggest that the absence of a comprehension benefit in older adults is linked to a change in the relationship between audiovisual intermodulation and behaviour, despite largely preserved task-sensitive brain responses.

## Introduction

In a crowded restaurant, a listener leans in, watches the speaker’s lips, and somehow pulls one voice out of the din. This everyday feat becomes markedly harder with age. Older adults often experience substantial declines in speech-in-noise perception, even in the absence of clinically significant hearing loss [1]. Importantly, compensatory strategies that were once effective - notably, watching the talker’s face - may no longer provide the same benefit. In these environments, successful speech comprehension depends on segregating a target speaker from competing streams and accurately aligning speech cues across auditory and visual modalities. However, ageing reduces temporal precision, weakens neural representations of speech, and impairs attentional control and working memory, making it more difficult to segregate competing speech streams and exploit predictive audiovisual cues [2–4]. These changes increase listening effort and contribute to poorer speech comprehension under adverse listening conditions [4,5]. Understanding how ageing affects the neural processes that support audiovisual speech perception is therefore essential for explaining age-related declines in speech comprehension.

Speech perception is inherently multisensory, relying on the integration of auditory and visual information, particularly in challenging listening environments. Visual speech cues, including lip movements and articulatory gestures, improve speech intelligibility by reducing ambiguity in the acoustic signal, especially when speech is masked by background noise [6] and competing speech [7,8]. Because visual speech typically precedes the corresponding acoustic signal, it provides predictive information that facilitates the processing of upcoming auditory input [9,10]. A key feature supporting this process is the speech amplitude envelope, which captures the slow temporal fluctuations underlying the rhythmic and syllabic structure of speech and is represented in both auditory and visual signals [11]. This shared temporal structure supports the alignment and integration of auditory and visual speech during perception [8,12].

Older adults continue to benefit from both auditory and visual speech, but growing evidence suggests that the neural mechanisms supporting audiovisual processing change with age. Compared to younger adults, older adults often rely more heavily on visual speech cues, particularly under adverse listening conditions [13,14]. This increased visual reliance has frequently been interpreted as a compensatory response to declining auditory reliability, whereby additional sensory or cognitive resources are recruited to maintain successful speech perception [9,15,16]. However, an alternative neurocognitive ageing account proposes that ageing reduces the functional specificity of neural responses, resulting in broader and less selective cortical representations, a process known as cortical dedifferentiation [17–19]. Age-related changes in audiovisual speech perception could therefore arise from changes within the auditory and visual systems themselves, resulting in less selective representations of speech information. Alternatively, sensory representations may remain relatively preserved, while the processes that combine auditory and visual information become altered with age. Although both accounts predict age-related changes in neural activity, they make different predictions about where in the processing hierarchy these changes arise. Distinguishing between them therefore requires methods that can separately assess modality-specific sensory responses and audiovisual interactions, while also determining how each relates to behavioural performance.

Previous neuroimaging and electrophysiological studies have begun to characterise both the sensory representations and broader cortical networks involved in audiovisual speech processing. Functional neuroimaging studies in young adults have consistently implicated superior temporal and auditory regions in audiovisual speech processing [20,21]. Electrophysiological studies have further shown that neural activity tracks the temporal dynamics of speech and is shaped by attention and predictive visual information [12,22–24]. In older adults, studies of healthy ageing have identified altered patterns of cortical recruitment during speech perception that are often interpreted as compensatory neural activity [25,26], and neural speech tracking also remains robust, although ageing has been associated with changes in both the magnitude and timing of speech-related neural responses, including reports of enhanced speech-envelope tracking despite poorer speech-in-noise perception [22,23]. Although these approaches reveal how auditory and visual speech signals are represented, they do not directly separate modality-specific sensory responses from the nonlinear neural interactions that emerge through their integration.

Rapid Invisible Frequency Tagging (RIFT) provides a direct way to investigate the neural mechanisms underlying audiovisual integration. Frequency tagging tracks sensory processing by modulating a stimulus at a specific frequency, eliciting a corresponding steady-state neural response [27,28]. RIFT extends conventional frequency tagging by using imperceptible high-frequency luminance modulation to measure visual processing without altering the perceptual experience of the stimulus [29]. In the present study, visual speech was frequency tagged using RIFT, while auditory speech was independently tagged at a different frequency, enabling modality-specific neural responses to be measured simultaneously [30–33]. Crucially, nonlinear interactions between the independently tagged auditory and visual signals generate new frequency components at the sum or difference of the primary tagged frequencies, known as intermodulation frequencies. These responses are absent from the stimulus itself and therefore provide a direct neural signature of audiovisual integration rather than independent sensory processing [30,31,34,35].

Using this Rapid Invisible Frequency Tagging approach, we recently investigated audiovisual speech processing in younger adults by embedding either a behaviourally relevant or irrelevant speech envelope within the RIFT signal applied to a speaker’s mouth region [36]. We showed that embedding the behaviourally relevant speech envelope improved speech comprehension. MEG recordings showed enhanced visual tagging responses, reduced auditory tagging responses, and stronger intermodulation responses when the visual RIFT conveyed behaviourally relevant temporal information. Stronger intermodulation responses in the left inferior frontal cortex predicted greater improvements in speech comprehension, linking nonlinear audiovisual integration to successful speech perception. Whether ageing preserves this coupling between behaviourally relevant audiovisual integration and successful speech comprehension has yet to be established.

To distinguish whether age-related changes in audiovisual speech perception arise primarily from altered sensory representations or from changes in the way auditory and visual information are integrated, we applied the RIFT paradigm to healthy older adults. Participants completed a dichotic listening task in which they attended to one of two competing speech streams while visual speech was frequency tagged at 55 Hz and auditory speech at 40 Hz. The visual tagging signal was modulated by either the attended or unattended speech envelope, allowing us to manipulate the behavioural relevance of visual temporal information while independently measuring auditory and visual sensory responses and their nonlinear interactions. Our primary behavioural hypothesis was that embedding the attended speech envelope within visual speech would improve speech comprehension, as previously observed in younger adults. At the neural level, we expected modality-specific sensory responses to remain sensitive to task relevance in older adults. Specifically, visual tagging responses were expected to increase when visual speech carried the attended speech envelope, whereas auditory tagging responses were expected to be stronger when visual speech carried the unattended speech envelope, consistent with our findings in younger adults. In contrast, if ageing primarily affects audiovisual integration rather than sensory encoding, we expected intermodulation responses to be reduced or otherwise altered in older adults. We further predicted that any age-related reduction in the behavioural benefit of Task-Relevant modulation would be associated with weaker intermodulation responses.

## Materials and Methods

The study design, hypotheses, and primary analysis plan were preregistered on the Open Science Framework prior to data analysis (OSF; 10.17605/OSF.IO/2P43X).

### Participants

46 healthy older adults (25 females; aged 65-78 years; mean age = 71 ± 3.8 years, mean ± SD) were recruited and tested at the Centre for Human Brain Health (CHBH), University of Birmingham. Data from 6 participants were excluded prior to analysis due to excessive recording noise, technical issues, or inadequate task performance. To ensure data quality, a minimum performance criterion was applied: individuals scoring below 50% accuracy on the comprehension task were removed. As the task used a three-alternative forced-choice design (chance level = 33.3%), this threshold ensured inclusion of only those performing reliably above chance. After applying these criteria, the final sample consisted of 40 participants (22 female; age range 65-78 years; mean age = 70 ± 3.6 years, mean ± SD). All participants reported no past or present neurological, psychiatric, or neurodevelopmental disorders, and none indicated the use of antipsychotic medication. Written informed consent was obtained from all individuals before participation, and financial compensation was provided. All participants were right-handed, as determined by the Edinburgh Handedness Inventory [37]. In addition, they were native speakers of English, with either normal or corrected-to-normal vision and normal auditory function. Hearing ability was assessed using the Etymotic Research at-home hearing test (Etymotic Research, Elk Grove Village, IL, USA), alongside a self-report measure of auditory function using the Speech, Spatial and Qualities of Hearing Scale [38]. This approach is supported by population-based evidence indicating that individuals without clinically relevant hearing impairment typically report SSQ scores in the upper range of the scale (approximately ≥7), with only modest effects of age [39]. All participants scored 23 or above on the Montreal Cognitive Assessment (MoCA; Nasreddine et al., 2005). This cutoff was chosen based on evidence that the original threshold of 26 may overestimate impairment, and that a lower cutoff of 23 may provide a more appropriate balance between sensitivity and specificity [41].

For comparison, a younger adult cohort (N = 40; 20 female; aged 19-35 years; mean age = 23.4 ± 2.9 years) was tested using the same experimental paradigm and comparable inclusion criteria (young adult data were previously analysed and reported on, Reynolds et al, (2026), addressing research questions distinct from those examined here. Ethical approval for the study was granted by the University of Birmingham STEM ethics committee (ERN_18-0226AP28A), and all procedures conformed to the Declaration of Helsinki.

### Frequency Tagging

Stimuli were presented using a PROPixx DLP LED projector (VPixx Technologies Inc., Canada), capable of achieving a refresh rate of up to 1440 Hz. This was accomplished by dividing each 120 Hz frame from the graphics card into four quadrants, effectively increasing the temporal resolution to 480 Hz. Visual frequency tagging was applied to a rectangular region (120 by 90 pixels) surrounding the speaker’s lip movements. This region was modulated using a 55 Hz sinusoidal luminance signal. A photodiode was positioned over a small square (30 by 30 pixels) in the lower-right corner of the display to record the timing of the visual flicker. Two tagging conditions were implemented: (1) constant-amplitude tagging and (2) amplitude-modulated tagging. In the constant condition, the 55 Hz luminance modulation was applied continuously at a fixed amplitude. In the amplitude-modulated conditions, the tagging signal was modulated over time by the amplitude envelope of the auditory speech signal (see *Stimuli, Task and Experimental Conditions* for details). This manipulation introduced slow, visible fluctuations in luminance. In all tagging conditions, the auditory signal was independently frequency-tagged at 40 Hz by modulating the speech waveform with a sinusoidal signal.

### Stimuli, Task, and Experimental Conditions

The stimuli consisted of continuous audiovisual speech recordings adapted from TED talks and produced by a professional male speaker. Each video lasted approximately 7-9 minutes and has been validated in previous work [8,36]. During each block, participants viewed the video while simultaneously listening to two different speech streams presented dichotically, with one stream delivered to each ear. At the beginning of each block, participants were directed which ear to attend to (task-relevant speech) and which to ignore (task-irrelevant speech). The attended ear remained constant across all conditions within participants but was counterbalanced across the sample (half attending left, half right). Participants were not informed that the attended ear did not change across conditions. The experiment comprised four conditions (Figure 1), of which three were included in the main analyses (see Supplementary Materials for additional conditions). In the task-relevant amplitude modulation condition (Task-Relevant AM), the visual tagging signal (55 Hz) was modulated by the amplitude envelope of the attended speech. In the task-irrelevant amplitude modulation condition (Task-Irrelevant AM), the same visual tagging signal was modulated by the envelope of the unattended speech. In the pure tagging condition (Pure Tagging), the visual tagging signal was applied at a constant amplitude without modulation. In all tagging conditions, the auditory speech signal was independently frequency-tagged at 40 Hz without amplitude modulation. A separate no-tagging condition (No Tagging), in which neither auditory nor visual tagging was applied, was included as a control. Although the 40 Hz auditory modulation was perceptible, accuracy was not reduced in the tagged conditions relative to No Tagging, and response times were faster, providing no evidence that the tagging procedure impaired speech comprehension (Supplementary Fig. 1).

**Fig. 1.**
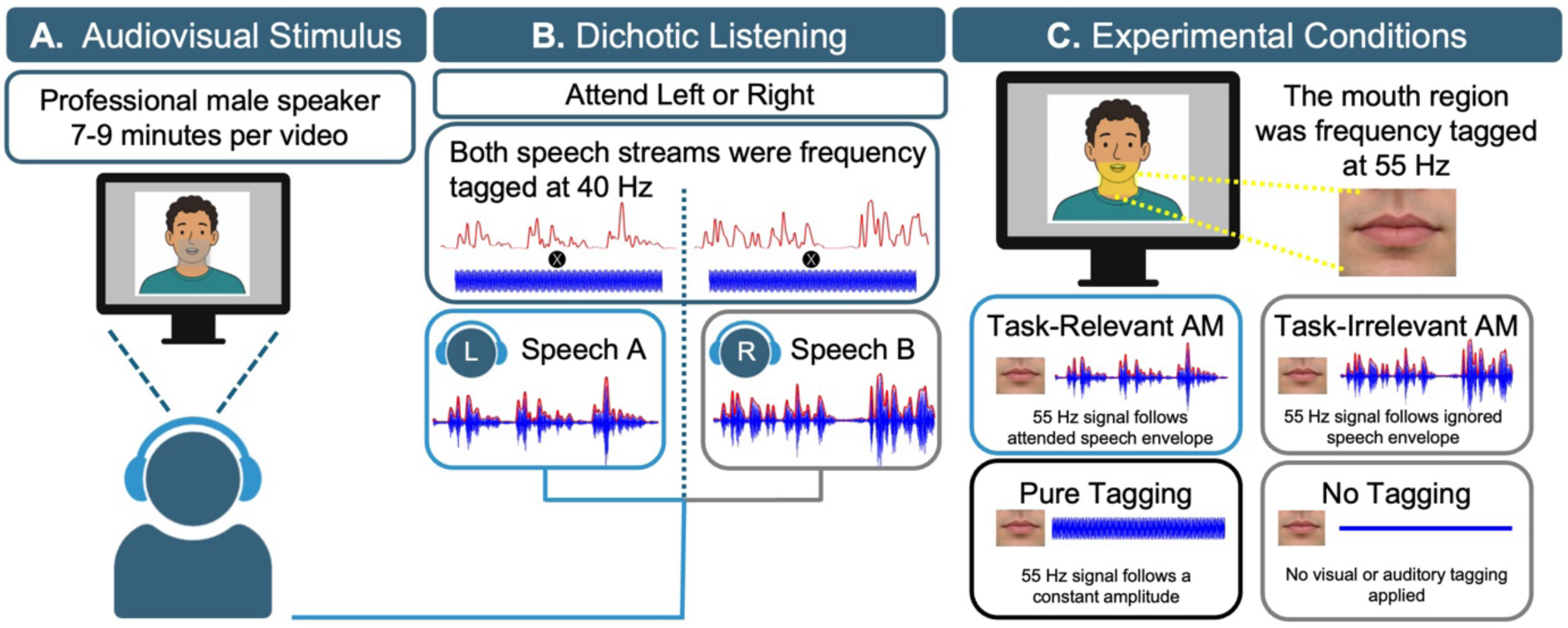
| Experimental paradigm and frequency tagging conditions. Computer-generated schematic representation of a speaker, included for illustrative purposes. The experimental stimuli consisted of naturalistic speech videos. **(A)** Participants viewed an audiovisual speech video. **(B)** Participants attended to one of two dichotic speech streams (blue line reflects attended speech, grey reflects ignored speech) which were frequency tagged at 40 Hz. **(C)** Visual tagging at 55 Hz (RIFT) was applied to the mouth region of the speaker, with its amplitude modulated by either the attended speech, unattended speech, or not at all.

### Experimental Procedure

Participants were seated comfortably in the MEG system, with the gantry positioned at 60° within a magnetically shielded room to minimise environmental interference. Stimuli were presented on a projection screen at a fixed viewing distance of 100 cm using Psychtoolbox [42] implemented in MATLAB (MathWorks, Natick, MA). Participants completed four audiovisual speech blocks, each corresponding to a different experimental condition. The order of conditions was randomised across participants. During each block, participants were instructed to attend to the designated auditory stream while viewing the video. Following each block, participants completed a 10-item multiple-choice questionnaire assessing comprehension of the attended speech. Both accuracy and response times were recorded. Each block, including the comprehension task, lasted approximately 10 minutes.

### Data Acquisition

Magnetoencephalography (MEG) data were recorded using a 306-channel Elekta Neuromag TRIUX system (MEGIN, Finland), comprising 102 magnetometers and 204 planar gradiometers. Signals were sampled at 1000 Hz and band-pass filtered online between 0.1 and 330 Hz. The visual tagging signal was monitored using a custom-built photodiode (Aalto Neuroimaging Centre, Finland) attached to the display, allowing precise recording of luminance modulation timing. Auditory stimuli were recorded directly from the MEG soundcard during presentation. Head position was tracked using four head position indicator (HPI) coils placed on the scalp (two on the forehead and one on each mastoid). The locations of the nasion, left and right preauricular points, and HPI coils were digitised using a Polhemus Fastrack system (Polhemus Inc., USA). In addition, approximately 300 scalp surface points were digitised to enable accurate co-registration with anatomical MRI data. Electrooculography (EOG) and electrocardiography (ECG) were recorded concurrently to monitor eye movements and cardiac activity. Behavioural responses were collected using a five-button response box (NAtA Technologies, USA).

### Eye-Tracking

Eye movements were recorded using the EyeLink 1000 Plus system (SR Research, Ottawa, Canada) at a sampling rate of 1000 Hz to ensure participants maintained fixation on the speaker’s face. A standard five-point calibration and validation procedure was performed at the start of each condition, and calibration was accepted only if all points were within 1° of visual angle. Eye-tracking data were successfully acquired for all younger adult participants. In the older adult group, eye-tracking data were not obtained for a subset of participants (N=11) due to calibration difficulties and data loss, which are common in ageing populations (Shoenfelt et al., 2025). Group-averaged gaze distributions for each experimental condition are shown in Supplementary Fig. S2.

### Data Preprocessing

MEG data were preprocessed using MNE-Python [44] following procedures adapted from the FLUX pipeline [45]. Noisy channels were identified and reconstructed prior to further processing. Signal Space Separation (SSS; Taulu and Kajola, (2005)) was applied to suppress external interference, and spatiotemporal SSS (tSSS; Taulu and Simola, (2006)) was used for datasets with elevated noise levels. Independent component analysis (ICA; Hyvärinen and Oja, (2000) was used to remove ocular and cardiac artefacts. The recordings were downsampled to 250 Hz. The continuous signal was then divided into 4-second epochs for analysis.

### MEG-MRI Co-Registration and Source-level Preprocessing

Structural MRI data were acquired for each participant using a T1-weighted magnetization-prepared rapid gradient-echo (MPRAGE) sequence (TR = 2000 ms, TE = 2.03 ms, TI = 880 ms, flip angle = 8°, field of view = 256 × 256 × 208 mm, voxel size = 1 mm isotropic). Cortical surface reconstruction was performed using FreeSurfer [49]. A boundary element model (BEM) was constructed using a watershed algorithm to define the scalp, inner skull, and outer skull surfaces [44]. Source space was defined on the reconstructed cortical surface, and forward solutions were computed for each participant [50,51]. Co-registration between MEG and MRI data was achieved by aligning digitised fiducial points (nasion and bilateral preauricular points) with the corresponding anatomical landmarks. This initial alignment was refined using an iterative closest point (ICP; Dale et al., (2000) algorithm to match the digitised head shape to the MRI-derived scalp surface [53]. Source estimates were computed using dynamic statistical parametric mapping (dSPM), allowing cortical activity to be projected onto a common anatomical space for group-level analysis.

### Behavioural Analysis

Behavioural performance was assessed using comprehension accuracy (%) and response time (RT) obtained from the speech-in-noise comprehension task. Analyses were conducted in Python. To examine condition effects within the older adult cohort, separate one-way repeated-measures analyses of variance (ANOVAs) were performed for accuracy and RT, with Condition (Task-Relevant AM, Task-Irrelevant AM and Pure Tagging) entered as a within-subject factor. Significant omnibus effects were followed by pairwise paired-samples t-tests. To control for multiple comparisons, Bonferroni-corrected p-values were used, with statistical significance defined as p < 0.05. Age-related differences in behavioural performance were examined using the three conditions (Task-Relevant, Task-Irrelevant, and Pure Tagging). Separate mixed-design ANOVAs were conducted for accuracy and RT, with Age Group (older adults, younger adults) entered as a between-subjects factor and Condition entered as a within-subject factor. Main effects of Age Group and Condition, as well as the Age Group by Condition interaction, were evaluated. Significant effects were followed by independent-samples t-tests comparing age groups within each matched condition. Resulting p-values were corrected for multiple comparisons using the Benjamini-Hochberg false discovery rate (FDR) procedure, with significance assessed at p < 0.05. Bonferroni correction was used for the small, prespecified set of within-group condition contrasts, whereas FDR correction was used for the broader family of between-group follow-up comparisons to control the false discovery rate while retaining statistical power.

### Frequency Tagging Analysis

To quantify tagging responses at the source level, coherence was computed between the seed channels and each cortical vertex using MNE-Python. The visual seed was defined by the photodiode signal recorded from the display, and the auditory seed was derived from the soundcard recordings of the presented speech signals. Coherence was estimated for each 4-second epoch using a multitaper spectral approach and averaged within ±1 Hz of the visual tagging frequency (55 Hz) and the auditory tagging frequency (40 Hz). Coherence values were averaged across epochs to obtain condition-specific source maps for each participant. These maps were subsequently morphed to a common template brain (FreeSurfer average) to enable group-level comparisons. Condition-related changes in tagging coherence were evaluated using non-parametric cluster-based permutation test. Individual participant coherence contrasts were first computed between the conditions of interest. Group-level inference was then carried out using one-sample tests against zero. Neighbouring source locations showing effects exceeding a predefined statistical threshold were combined into spatial clusters based on their adjacency structure. The strength of each cluster was quantified by the summed t-statistic across all constituent locations. A reference distribution was generated from 1,024 random sign-flip permutations, allowing cluster-level significance to be assessed while accounting for multiple comparisons. Observed clusters were deemed significant when their permutation-derived p-value was less than 0.05 (two-tailed).

### Intermodulation Frequency Analysis

To investigate neural integration of auditory and visual frequency-tagged inputs, intermodulation reference signals were constructed from the recorded auditory and photodiode channels. The auditory (40 Hz) and visual (55 Hz) signals were first isolated using narrow band-pass filters (±3 Hz around each tagging frequency). Sinusoidal models were then fitted to the filtered signals, and the resulting waveforms were multiplied to generate an intermodulation reference signal. This signal was subsequently normalised and filtered within the 1-20 Hz range. For each participant and experimental condition, coherence was estimated between the intermodulation reference and source-reconstructed cortical activity. Intermodulation responses were quantified at the 15 Hz difference frequency (55 − 40 Hz) by averaging coherence estimates within ±1 Hz of 15 Hz. Individual coherence maps were transformed to the FreeSurfer average template to enable group-level analyses. Differences between conditions were assessed using cluster-based permutation statistics to identify cortical regions showing reliable modulation of intermodulation coherence. Group-level inference was performed by computing condition contrasts for each participant at every cortical vertex and evaluating these values against zero using non-parametric one-sample permutation testing. Spatially neighbouring vertices were grouped according to the FreeSurfer average source-space adjacency structure. Candidate clusters were identified using a threshold derived from the two-tailed t-distribution (α = 0.05), and their significance was determined from a null distribution generated using 1,024 sign-flip permutations. Clusters were considered statistically significant when the resulting cluster-level p-value was below 0.05. To examine whether condition-related changes in intermodulation coherence were associated with response time, the Task-Relevant AM minus Task-Irrelevant AM difference in 15 Hz coherence was correlated with the corresponding difference in response time across participants. Spearman correlations were calculated at each cortical vertex and assessed using the same cluster-based permutation framework used for the accuracy analysis.

### Age Comparison Analysis

To examine age-related differences in source-level coherence, analyses were performed on the visual (55 Hz photodiode), auditory (40 Hz soundcard), and intermodulation (15 Hz) coherence maps using the three experimental conditions common to both age groups (Task-Relevant AM, Task-Irrelevant AM, and Pure Tagging). Individual coherence maps were transformed to the FreeSurfer average template to enable group-level comparisons. The main effect of Age Group was assessed by first averaging coherence across the three conditions for each participant and comparing younger and older adults using non-parametric independent-samples cluster-based permutation tests. The main effect of Condition was evaluated by pooling participants across age groups and performing a non-parametric cluster-based repeated-measures permutation analysis. Significant condition effects were followed up using paired cluster-based permutation tests for each pairwise condition contrast (Task-Relevant AM vs Task-Irrelevant AM, Task-Relevant AM vs Pure Tagging, and Task-Irrelevant AM vs Pure Tagging), with Bonferroni correction applied across the three comparisons (α = 0.0167). To investigate the Age Group by Condition interactions, condition-difference maps were first computed within each participant (Task-Relevant AM vs Task-Irrelevant AM, Task-Relevant AM vs Pure Tagging, and Task-Irrelevant AM vs Pure Tagging) and subsequently compared between younger and older adults using non-parametric independent-samples cluster-based permutation tests. For all analyses, spatially neighbouring vertices were grouped according to the FreeSurfer average source-space adjacency structure. Candidate clusters were identified using thresholds derived from the appropriate theoretical test statistic (t- or F-distribution; α = 0.05), and their significance was determined from a null distribution generated using 1,024 random permutations. Clusters were considered statistically significant at a cluster-level threshold of p < 0.05, except for the three pairwise condition contrasts and the three Age Group by Condition contrasts, for which Bonferroni correction was applied across comparisons (α = 0.0167).

Finally, to examine whether condition-related changes in intermodulation coherence were associated with comprehension benefit, we calculated the Task-Relevant AM minus Task-Irrelevant AM difference in 15 Hz coherence at each cortical vertex and the corresponding difference in percentage accuracy for each participant. Spearman correlations were computed separately in younger and older adults, and the correlation maps were compared directly between age groups. Correlation coefficients were Fisher-transformed for cluster formation. Within each age group, accuracy differences were shuffled across participants; for the between-group comparison, group labels were shuffled while each participant’s coherence map and accuracy difference remained paired. Spatial clusters were identified using a two-tailed cluster-forming threshold of p < 0.01. Cluster significance was assessed using 1,000 permutations and a cluster-level threshold of p < 0.05, with the maximum cluster mass across both signs and the whole cortex used to account for multiple comparisons within each analysis.

## Results

To determine whether speech-envelope-modulated RIFT influences comprehension, we manipulated experimental conditions where visual frequency tagging is modulated by attended or unattended speech envelope. Participants listened to continuous audiovisual speech while selectively attending to one of two competing auditory streams. Behavioural performance was assessed using comprehension accuracy and response time following each experimental condition. Performance was compared across conditions in which the visual tagging signal was modulated by the attended speech stream (Task-Relevant AM), the unattended speech stream (Task-Irrelevant AM), or presented as a constant-frequency tagging signal without amplitude modulation (Pure Tagging). This enabled us to determine whether speech comprehension was influenced by the behavioural relevance of the speech envelope embedded within the visual tagging signal.

### Age-Related Differences in Comprehension Accuracy Are Specific to the Task-Relevant AM Condition

To investigate whether the behavioural effects of speech-envelope-modulated frequency tagging differed with age, performance in the older adult (OA) cohort was compared with that of a younger adult (YA) cohort across the Task-Relevant AM, Task-Irrelevant AM, and Pure Tagging conditions (Fig. 2A-B). For speech comprehension accuracy, a mixed-design ANOVA revealed that there was no significant main effect of Age Group, F(1,78) = 3.42, p = 0.068, ηp² = 0.042, and no significant main effect of Condition, F(2,156) = 2.05, p = 0.132, ηp² = 0.026. However, a significant Age Group by Condition interaction was observed, F(2,156) = 8.23, p < 0.001, ηp² = 0.095. Follow-up comparisons revealed significantly higher accuracy in YA relative to OA in the Task-Relevant AM condition (p < 0.001), whereas no significant age-group differences were observed in the Task-Irrelevant AM or Pure Tagging conditions. Response time was defined as the interval between the onset of the comprehension question and the participant’s button-press response, providing a post-stimulus measure of the speed of comprehension-related retrieval and decision-making. For response time, a mixed-design ANOVA revealed a significant main effect of Age Group, F(1,78) = 17.19, p < 0.001, ηp² = 0.181, indicating that OA responded more slowly than YA overall. A significant main effect of Condition was also observed, F(2,156) = 34.97, p < 0.001, ηp² = 0.310, indicating that RT varied across experimental conditions. Importantly, there was a significant Age Group by Condition interaction, F(2,156) = 4.28, p = 0.015, ηp² = 0.052, indicating that the magnitude of the age-related slowing differed across conditions. Follow-up t-tests with false discovery rate (FDR) correction showed significantly slower RTs in OA compared to YA in the Task-Relevant AM (p < 0.001), Task-Irrelevant AM (p < 0.001), and Pure Tagging (p = 0.013) conditions. These findings indicate that older adults showed slower behavioural responses across all conditions, whereas lower comprehension accuracy relative to younger adults was specific to the Task-Relevant AM condition.

**Fig. 2.**
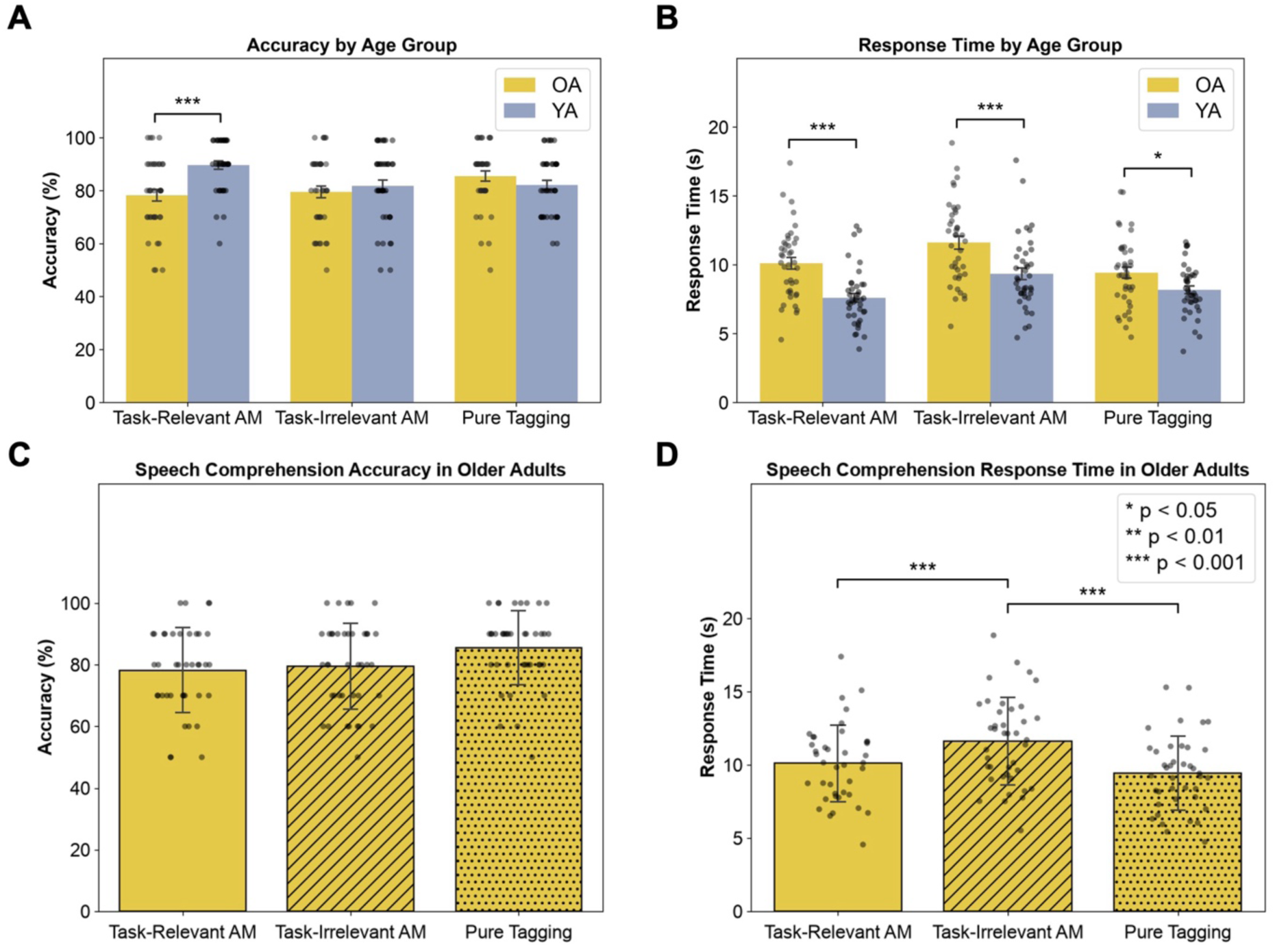
| Behavioural performance across experimental conditions in older and younger adults. **(A)** Accuracy comparison between younger adults (YA) and older adults (OA) for the matched experimental conditions. **(B)** Response time comparison between YA and OA for the matched experimental conditions. Individual participant data are overlaid as points. Significance markers in (A-B) indicate pairwise comparisons following Benjamini–Hochberg false discovery rate (FDR) correction for multiple comparisons. **(C)** Mean comprehension accuracy (%) in the older adult (OA) cohort across the Task-Relevant amplitude modulation (AM), Task-Irrelevant AM, and Pure Tagging conditions. **(D)** Mean response times (RT; s) for the OA cohort across the same conditions. Error bars represent ±1 SD. Significance markers in (C-D) indicate Bonferroni-corrected post hoc comparisons (p < 0.05, p < 0.01, p < 0.001).

#### Behavioural Effects of Speech-Envelope-Modulated Tagging in Older Adults

To determine whether speech-envelope-modulated frequency tagging influenced behavioural performance, comprehension accuracy and response time were examined across the Task-Relevant AM, Task-Irrelevant AM, and Pure Tagging conditions in older adults (Fig. 2C-D). Although the one-way repeated-measures ANOVA revealed a significant effect of condition on accuracy, F(2,78) = 4.19, p = 0.019, none of the Bonferroni-corrected pairwise comparisons reached significance. This indicates a condition-level effect on accuracy, but the differences were distributed across conditions rather than being driven by a single robust pairwise contrast. In contrast, response time varied significantly as a function of condition, F(2,78) = 22.33, p < 0.001. Follow-up pairwise comparisons revealed significantly slower responses in the Task-Irrelevant AM condition relative to both the Task-Relevant AM condition (p < 0.001) and the Pure Tagging condition (p < 0.001). No significant response time difference was observed between the Task-Relevant AM and Pure Tagging conditions (p = 0.108). Older adults responded more slowly than younger adults across conditions. Importantly, this general slowing did not mirror the condition-specific accuracy difference, arguing against a simple speed-accuracy trade-off as an explanation for the reduced Task-Relevant accuracy benefit in older adults.

#### Older Adults Show Greater Auditory Tagging Coherence, with Comparable Visual and Intermodulation Responses Across the Groups

To determine whether ageing altered overall neural responses to frequency-tagged audiovisual speech, source-level coherence maps were first averaged across the three shared experimental conditions (Task-Relevant AM, Task-Irrelevant AM, and Pure Tagging) within older and younger adults group and compared between them using cluster-based permutation testing. No significant age-related differences were observed for visual tagging coherence at 55 Hz (photodiode; all cluster-corrected p > 0.05) or for intermodulation coherence at 15 Hz (all cluster-corrected p > 0.05), with no evidence of an overall age-group difference in visual tagging or intermodulation responses when averaged across conditions. In contrast, a significant Age Group difference was observed for auditory tagging coherence at 40 Hz, with older adults exhibiting greater coherence than younger adults within a single significant temporal cluster (cluster-corrected p = 0.018; Fig. 3A). These findings indicate that ageing was associated with greater coherence between brain activity and the auditory tagging signal, whereas no significant age-group differences were observed in visual tagging or intermodulation responses. Mean coherence values extracted from the significant cluster (Fig. 3B) showed a similar pattern of age-related differences across the three experimental conditions. No significant age-group differences were observed for any of the three condition contrasts in visual, auditory, or intermodulation coherence (all cluster-corrected p > 0.0167), providing no evidence that condition-dependent modulation differed between age groups.

**Fig. 3.**
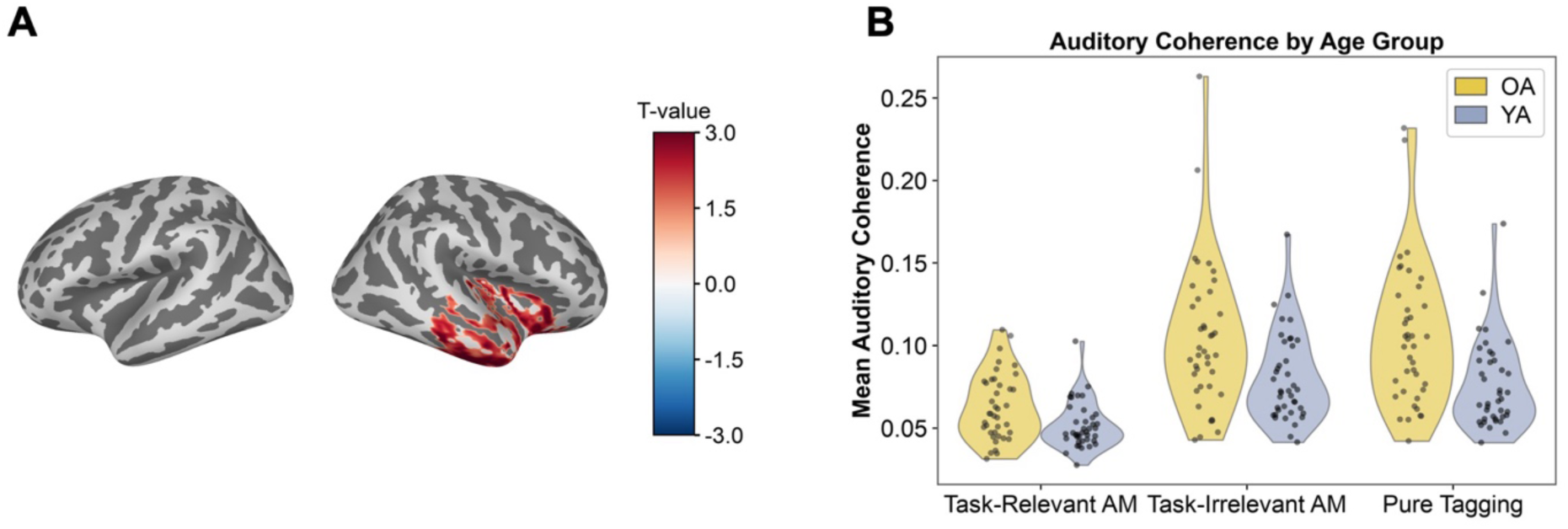
| Age-related increase in auditory tagging coherence. **(A)** Whole-brain cluster-based permutation analysis identified a significant right temporal cluster in which auditory 40 Hz coherence was greater in older adults than younger adults (cluster-corrected *p* = 0.018). Brain maps display *T*-values on the inflated FreeSurfer average cortical surface. **(B)** Mean coherence values extracted from the significant cluster for each condition and age group. Older adults showed greater auditory coherence than younger adults, with a similar pattern of group differences across all three conditions. No significant interaction between age group and condition was observed, indicating that the magnitude of the age effect did not differ between Task-Relevant AM, Task-Irrelevant AM, and Pure Tagging conditions.

#### Condition Effects on Neural Tagging Are Comparable Across Age Groups

To determine how speech-envelope modulation influenced neural tagging responses across age groups, source-level coherence was compared across the Task-Relevant AM, Task-Irrelevant AM, and Pure Tagging conditions using data pooled across younger and older adults (Fig. 4). Significant main effects of condition were observed for visual tagging coherence at 55 Hz, auditory tagging coherence at 40 Hz, and intermodulation coherence at 15 Hz, and were followed by Bonferroni-corrected pairwise cluster-based permutation tests. Across measures, the pattern showed that neural responses remained sensitive to whether the visual modulation carried the attended or unattended speech envelope. Visual coherence was greater when the tagging signal tracked the attended rather than unattended speech stream (Fig. 4A; all p ≤ 0.0127), while auditory coherence showed the opposite pattern, with reduced responses in the Task-Relevant AM condition relative to the Task-Irrelevant AM (Fig. 4D; both p < 0.001) and Pure Tagging (Fig. 4E; both p < 0.001) conditions. Visual coherence also differed from Pure Tagging in both the Task-Relevant (Fig. 4B; both p = 0.0010) and Task-Irrelevant (Fig. 4C; both p = 0.0010) AM conditions. Intermodulation coherence was likewise greater in the Task-Relevant than Task-Irrelevant AM condition (Fig. 4F; both p < 0.001). Crucially, none of these condition effects differed significantly between younger and older adults, with no Age Group by Condition interactions for visual, auditory, or intermodulation coherence (all cluster-corrected p > 0.05). Overall, condition-dependent changes in visual, auditory, and intermodulation coherence were comparable across age groups.

**Fig. 4.**
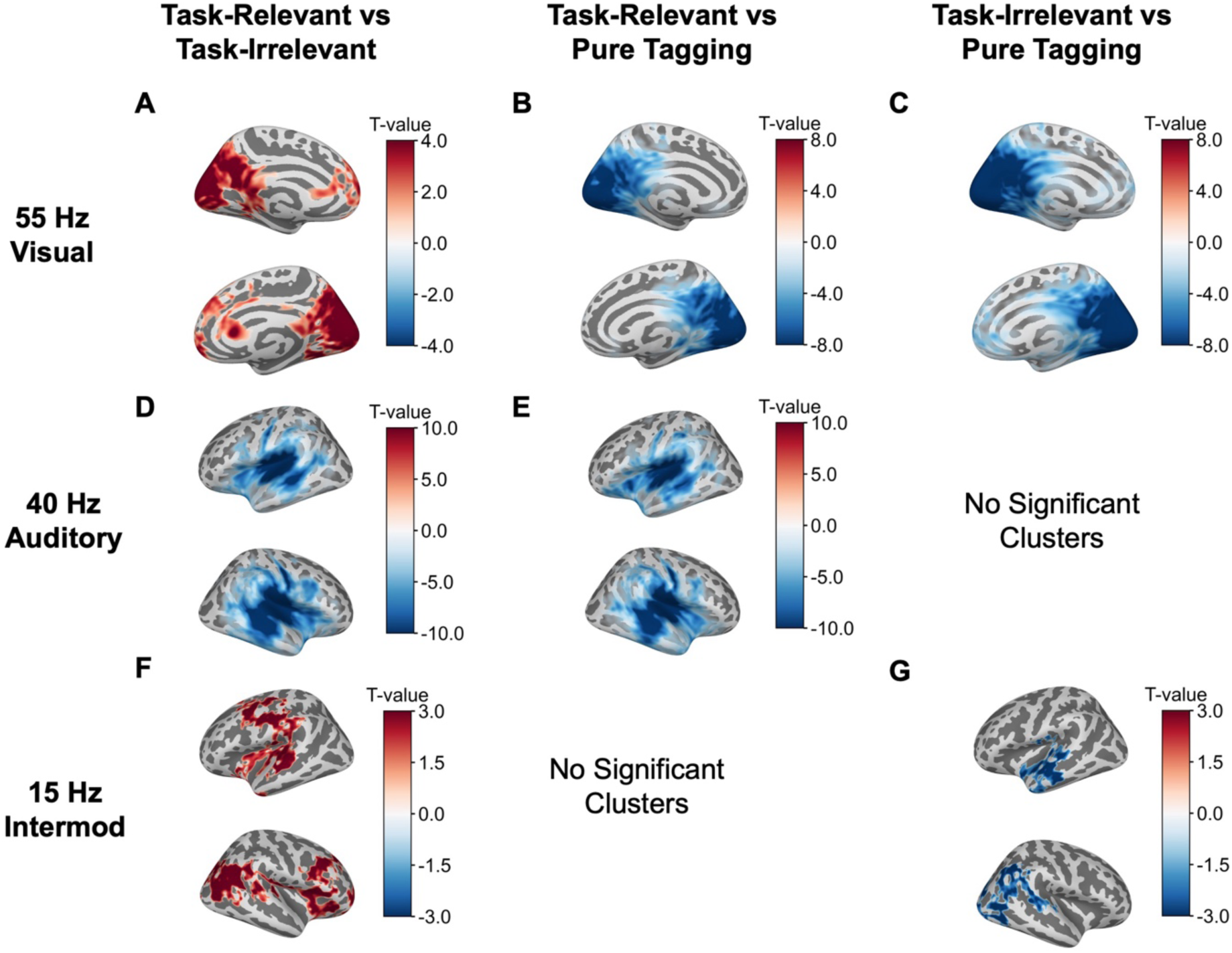
| Condition effects on neural tagging responses across age groups. (A-C) Source-level differences in visual (55 Hz) coherence for the Task-Relevant amplitude modulation (AM) versus Task-Irrelevant AM **(A)**, Task-Relevant AM versus Pure Tagging **(B)**, and Task-Irrelevant AM versus Pure Tagging **(C)** comparisons. **(D-E)** Corresponding source-level differences in attended auditory (40 Hz) coherence. **(F-G)** Corresponding source-level differences in audiovisual intermodulation (15 Hz) coherence. Data were pooled across younger and older adults to assess the main effect of condition. Warm colours indicate greater coherence in the first condition of each comparison, whereas cool colours indicate greater coherence in the second condition. Only clusters surviving Bonferroni correction across the three pairwise condition contrasts (α = 0.0167) are displayed.

#### Condition Effects on Neural Tagging in Older Adults Mirror Those in Younger Adults

To determine how speech-envelope modulation affected neural tagging in older adults, source-level coherence was compared between the Task-Relevant AM, Task-Irrelevant AM and Pure Tagging conditions (Fig. 5). Visual tagging was sensitive to the behavioural relevance of the modulating speech envelope: coherence at 55 Hz was significantly greater in the Task-Relevant AM than the Task-Irrelevant AM condition across four occipital clusters (Fig. 5A; all p < 0.05). However, coherence in both amplitude-modulated conditions was lower than during Pure Tagging, with significant reductions for Task-Relevant AM across three occipital clusters (Fig. 5B; all p < 0.05) and for Task-Irrelevant AM across two occipital clusters (Fig. 5C; all p < 0.05). Thus, while the cleaner, unmodulated 55-Hz signal produced the strongest overall visual tagging response, modulation by the attended rather than unattended speech envelope enhanced visual coherence. Auditory tagging showed a different pattern. At 40 Hz, coherence was significantly lower in the Task-Relevant AM than the Task-Irrelevant AM condition across two bilateral temporal clusters (Fig. 5D; both p < 0.001) and was also lower than in the Pure Tagging condition across two temporal clusters (Fig. 5E; both p < 0.001). Task-Irrelevant AM and Pure Tagging did not differ significantly, indicating that the reduction in auditory coherence was specific to the Task-Relevant condition rather than a general consequence of amplitude modulation. Critically, audiovisual intermodulation showed the opposite task-relevance effect: 15 Hz coherence was significantly greater in the Task-Relevant AM than the Task-Irrelevant AM condition across three bilateral clusters spanning sensorimotor and frontoparietal regions, with additional right temporal and occipital involvement (Fig. 5F; all p < 0.05). Task-Relevant AM did not differ significantly from Pure Tagging, whereas Task-Irrelevant AM showed lower coherence than Pure Tagging within one cluster (Fig. 5G; p < 0.005). Together, these results show that older adults remained sensitive to which speech stream modulated the visual signal, with task relevance affecting visual, auditory and audiovisual responses. This overall pattern was broadly consistent with that previously observed in younger adults using the same paradigm [36].

**Fig. 5.**
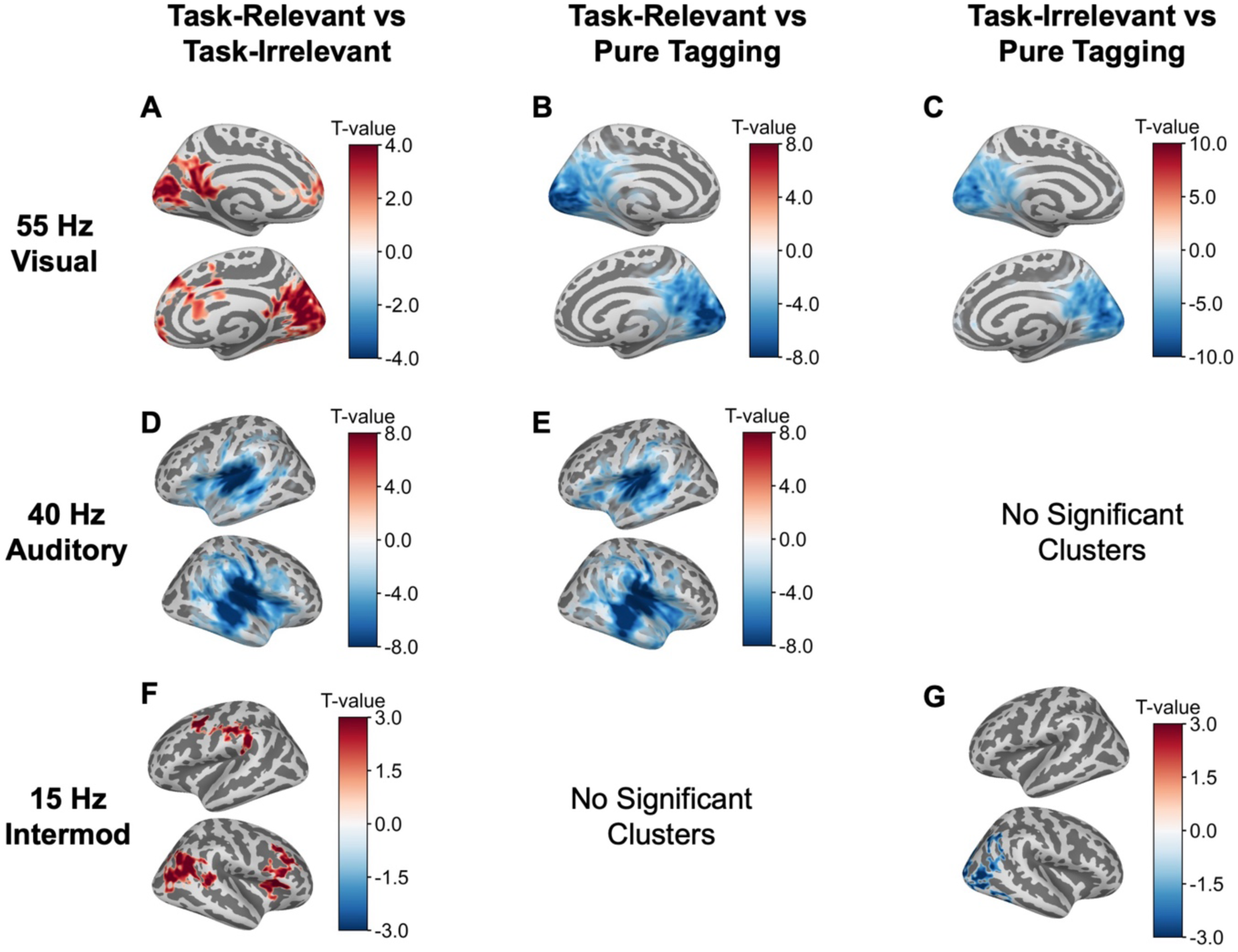
| Within-group (older adult) condition differences in coherence. (A-C) Source-level differences in visual (55 Hz) coherence for the Task-Relevant amplitude modulation (AM) versus Task-Irrelevant AM **(A)**, Task-Relevant AM versus Pure Tagging **(B)**, and Task-Irrelevant AM versus Pure Tagging **(C)** comparisons. **(D-E)** Source-level differences in attended auditory (40 Hz) coherence for the Task-Relevant AM versus Task-Irrelevant AM **(D)** and Task-Relevant AM versus Pure Tagging **(E)** comparisons. **(F-G)** Source-level differences in audiovisual intermodulation (15 Hz) coherence for the Task-Relevant AM versus Task-Irrelevant AM **(F)** and Task-Irrelevant AM versus Pure Tagging **(G)** comparisons. Warm colours indicate greater coherence in the first condition of each comparison, whereas cool colours indicate greater coherence in the second condition. Only vertices belonging to significant clusters identified using cluster-based permutation testing (p < 0.05, corrected) are displayed. No significant clusters were observed for the Task-Irrelevant AM versus Pure Tagging comparison at 40 Hz or the Task-Relevant AM versus Pure Tagging comparison at 15 Hz.

#### The Relationship Between Task-Relevant Intermodulation Responses and Comprehension Is Weaker in Older Adults

To determine whether changes in audiovisual intermodulation were related to the comprehension benefit of task-relevant speech-envelope modulation, we correlated the Task-Relevant minus Task-Irrelevant difference in 15 Hz intermodulation coherence with the corresponding difference in accuracy (Fig. 6). In younger adults, a significant positive cluster was identified in left temporal cortex (Fig. 6A-B; p = 0.029). Participants who showed a greater increase in intermodulation coherence in the Task-Relevant condition also tended to show a greater improvement in comprehension accuracy. No significant clusters were identified within the older adult group. We then directly compared the correlation maps between age groups and identified a significant cluster in which the relationship between intermodulation coherence and accuracy differed (Fig. 6C-D; p = 0.035). This relationship was more positive in younger than in older adults, suggesting an age-related difference in how changes in audiovisual intermodulation relate to the comprehension benefit of task-relevant modulation. These findings indicate that although task-relevant modulation increased intermodulation coherence across age groups, individual differences in this response were linked to the comprehension benefit in younger adults, but this relationship differed in older adults. No significant source-level clusters were identified for the relationship between intermodulation coherence and response-time differences.

**Fig. 6.**
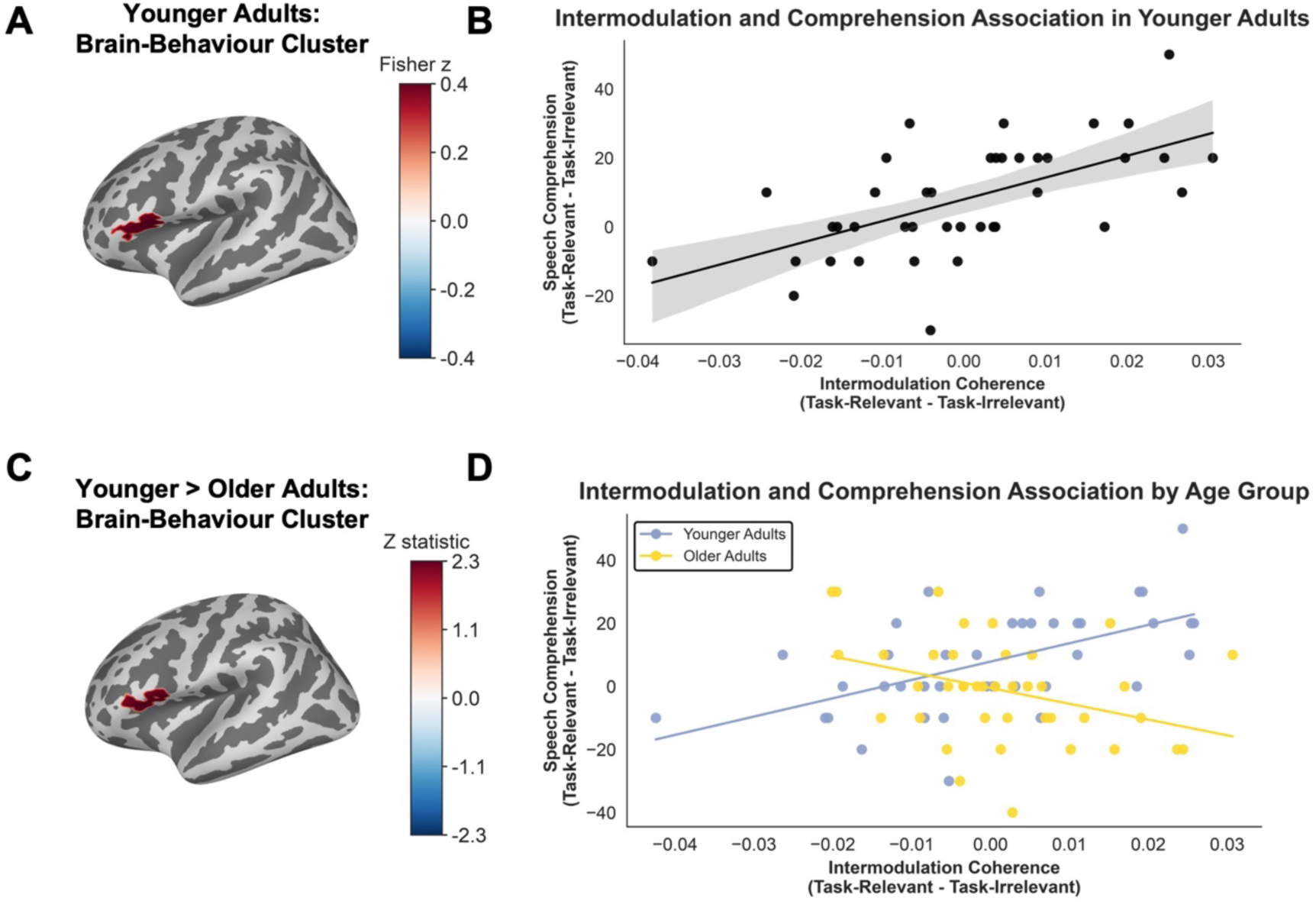
| Audiovisual intermodulation relates to speech comprehension benefit differently across age groups. **(A)** Source-level Spearman correlation between the Task-Relevant minus Task-Irrelevant difference in 15 Hz intermodulation coherence and the corresponding difference in comprehension accuracy in younger adults. The colour scale shows Fisher-transformed correlation coefficients. **(B)** Scatter plot of the two differences, averaged across the significant cluster in A. **(C)** Direct comparison of the younger- and older-adult correlation maps. Positive Z statistics indicate a stronger positive association in younger adults. **(D)** Scatter plot for the significant cluster in C, with younger adults in blue and older adults in yellow. Brain maps show clusters surviving spatial cluster correction at *p* < 0.05 within each analysis.

## Discussion

The present study investigated whether age-related declines in challenging listening conditions arise from impaired sensory encoding of audiovisual speech or from alterations in the neural processes that support audiovisual integration. Our findings do not provide evidence for a general decline in sensory encoding. Instead, age-related differences emerged primarily in the relationship between audiovisual integration and successful comprehension. Older adults failed to show the accuracy benefit from embedding the behaviourally relevant speech envelope within the RIFT visual tagging signal previously observed in younger adults (Fig. 2A). Nevertheless, older adults remained sensitive to the behavioural relevance of this information: visual and intermodulation coherence were enhanced when the visual modulation tracked the attended rather than unattended speech envelope. Crucially, these condition-dependent effects did not differ significantly between age groups. Instead, age-related differences emerged in the relationship between changes in audiovisual intermodulation and successful speech comprehension: whereas younger adults who showed a larger Task-Relevant increase in intermodulation also showed a larger comprehension benefit, this relationship was absent in older adults. Together, these findings suggest that age-related difficulties understanding speech in noisy environments may arise not from a failure to encode or integrate behaviourally relevant audiovisual information, but from a change in how these neural responses relate to successful comprehension.

This dissociation between preserved neural sensitivity and altered behavioural relevance has implications for theoretical accounts of age-related cognitive changes. The preserved condition-dependent modulation of visual, auditory, and intermodulation responses (Figs. 4 and 5) argues against a straightforward dedifferentiation account in which ageing results in broadly less selective neural representations [17–19]. Older adults continued to distinguish behaviourally relevant from irrelevant information across both modality-specific and audiovisual responses. At the same time, the enhanced auditory response in older adults (Fig. 3) could be consistent with compensatory accounts, whereby additional neural resources are recruited in response to age-related declines in processing efficiency [15,16]. However, because greater auditory coherence was not accompanied by better comprehension, this increased response cannot readily be interpreted as successful compensation. This is consistent with the Scaffolding Theory of Aging and Cognition, which proposes that additional neural recruitment may occur in response to age-related neural challenge without necessarily being sufficient to maintain performance [54]. This highlights an important distinction between preserved neural responses and the extent to which those responses support successful speech comprehension. This distinction was most evident in the relationship between audiovisual intermodulation and comprehension, where the clearest age-group difference emerged. Older adults continued to generate nonlinear audiovisual interactions that were sensitive to behaviourally relevant visual information. However, unlike younger adults in our previous study [36], older adults did not show a significant improvement in comprehension accuracy in the Task-Relevant relative to the Task-Irrelevant condition (Fig. 2C). In younger adults, greater Task-Relevant relative to Task-Irrelevant changes in intermodulation coherence were associated with a greater corresponding benefit in comprehension accuracy, with this relationship localised to left temporal cortex (Fig. 6A-B). Temporal regions form a central part of the cortical network supporting speech and language processing, including the mapping of speech signals onto linguistic representations [55,56]. Recent evidence further shows aligned auditory and visual phonological representations in posterior superior temporal and somatomotor cortex [57], suggesting that the present brain-behaviour relationship may involve regions in which heard and seen speech converge onto shared phonological representations. No significant brain-behaviour relationship was observed in older adults, and importantly, the relationship between intermodulation coherence and comprehension differed significantly between age groups within a left temporal cluster (Fig. 6C-D). Although audiovisual integration remained sensitive to task relevance in older adults, individual differences in these responses were not related to successful speech comprehension in the same way as in younger adults. This suggests that ageing may alter how audiovisual information supports speech comprehension, rather than audiovisual integration itself. Similar patterns have been reported in older adults, where multisensory processing can remain preserved or enhanced despite age-related differences in behavioural performance [58–60].

Previous studies have shown that visual speech continues to support speech perception in older adults, particularly under adverse listening conditions, with evidence for an increased reliance on visual information with age [13,14]. Our findings, however, point to a more specific age-related difference in the ability to exploit the temporal information provided by visual speech. Visual speech provides predictive information about the timing of upcoming auditory input, which can facilitate auditory processing and speech comprehension [9,10]. In the present paradigm, embedding the attended speech envelope within the visual modulation provided additional temporal information that was behaviourally relevant to the speech stream participants were required to follow. Although this did not produce the corresponding accuracy benefit in older adults, response times were slower following the Task-Irrelevant AM condition than following either the Task-Relevant AM or Pure Tagging conditions. This suggests that older adults were behaviourally sensitive to whether the visual modulation carried information from the attended or competing speech stream. However, because response times were measured during the subsequent comprehension questions rather than during speech presentation, they cannot be interpreted as a direct measure of online speech processing or listening effort. The longer responses following Task-Irrelevant modulation may instead be consistent with greater processing demands, as response latencies can increase under greater perceptual and cognitive demands [4,61]. Although older adults showed a clear response-time cost in the Task-Irrelevant condition, we did not identify a corresponding source-level relationship with intermodulation coherence. High accuracy may have made individual differences in performance harder to detect. Future studies could use more graded task difficulty and measures that place greater emphasis on response speed to better assess brain-behaviour relationships.

In contrast to the age-related differences in behaviour and brain–behaviour relationships, sensory responses to the auditory and visual speech signals were largely preserved in older adults. Visual tagging effects were centred in occipital cortex, while auditory tagging effects were distributed across temporal regions (Figs. 4 and 5), consistent with modality-specific sensory processing. Visual and auditory tagging responses remained sensitive to the behavioural relevance of the visual modulation, with no evidence that these condition-dependent effects differed between age groups. In particular, older adults showed enhanced visual tagging when the visual modulation tracked the attended rather than the unattended speech envelope, demonstrating that the visual system remained sensitive to the behavioural relevance of the embedded speech information. Auditory tagging showed the opposite pattern, with reduced coherence when the visual modulation carried the attended speech envelope. Rather than reflecting reduced auditory processing, this opposing pattern of enhanced visual and reduced auditory tagging may indicate a task-dependent reweighting of audiovisual speech processing, whereby behaviourally relevant visual information is weighted more strongly when it carries the temporal structure of the attended speech stream. Such reweighting is consistent with evidence that attention and task demands shape multisensory integration [62] and that visual speech can modulate auditory cortical processing [63].

This preservation extended beyond modality-specific responses to their interaction. Intermodulation responses at 15 Hz, reflecting nonlinear interactions between the auditory and visual tagging signals, also remained sensitive to the behavioural relevance of the visual modulation in older adults. Task-Relevant AM produced greater intermodulation coherence than Task-Irrelevant AM (Figs. 4F and 5F), and these condition-dependent effects did not differ significantly between age groups. The Task-Relevant effect was distributed across bilateral sensorimotor and frontoparietal regions, with additional right temporal and occipital involvement. The distribution of this effect suggests that when the visual envelope matched the attended speech stream, audiovisual interaction was strengthened across a broader speech-processing network rather than within sensory cortex alone. Temporal and occipital involvement is consistent with auditory and visual speech processing, while sensorimotor regions are thought to support the mapping between perceived speech and articulatory representations [56,64]. Recent evidence for shared auditory-visual phonological representations in somatomotor cortex further suggests that these regions may support interactions between auditory and visual speech at a phonological or articulatory level [57]. Frontoparietal involvement may additionally reflect cognitive-control processes that maintain and prioritise the behaviourally relevant speech stream over extended periods, consistent with the role of cingulo-opercular systems in sustained task control [65]. The stronger Task-Relevant intermodulation response may therefore reflect greater integration of visual timing information with the attended auditory speech stream across sensory, articulatory and attentional systems. This suggests that older adults still integrate behaviourally relevant audiovisual speech, but these neural responses are less closely linked to successful comprehension.

A key strength of the present study is the use of RIFT to dissociate modality-specific sensory responses from nonlinear audiovisual interactions within the same naturalistic speech paradigm. Several limitations should nevertheless be considered. First, the modest sample size limits our ability to capture the full extent of individual variability in behavioural and neural responses, particularly given the substantial heterogeneity reported across older adult populations [66]. Second, participants were healthy older adults selected to meet predefined hearing and cognitive criteria, which may limit generalisability to the broader ageing population. Finally, although intermodulation coherence provides a frequency-specific marker of nonlinear interaction between auditory and visual signals, it does not reveal the precise neural computations underlying this interaction. Future studies using larger and more heterogeneous samples will be important for determining which sensory and cognitive factors influence whether audiovisual brain responses support successful speech comprehension in later life.

## Conclusion

We found that age-related difficulties in speech perception under challenging listening conditions do not reflect a general decline in the neural encoding of auditory or visual speech. Healthy ageing did not abolish the neural signatures of audiovisual speech processing. Older adults’ visual, auditory and intermodulation responses remained sensitive to whether the visual signal carried the attended speech envelope, What differed was the behavioural payoff: older adults did not show the comprehension benefit seen in younger adults, and individual differences in audiovisual intermodulation did not track comprehension benefit as they did in younger adults. Age-related speech-in-noise difficulties may therefore arise less from how audiovisual speech is encoded and integrated than from how integrated information is used to support comprehension. Understanding why these preserved neural responses relate differently to comprehension in older adults will be important for explaining communication difficulties in challenging listening environments.

## Author Contributions

CR, KS, and HP conceived the study. KS and HP supervised the project. CR collected the data, analysed the data, and drafted the manuscript. All authors contributed to the interpretation of the results, revised the manuscript, and approved the final version.

## Acknowledgements

We thank the Centre for Human Brain Health (CHBH), University of Birmingham, for infrastructure and technical support. We are grateful to Jonathan Winter for MEG technical support and assistance with data acquisition. We also thank all participants for their time and engagement.

## Funding

CR was funded by a University of Birmingham College of Life and Environmental Sciences Studentship. KS was funded by BBSRC (UKRI 3572)

## Competing Interests

The authors declare no competing interests.

## Code Availability

The code used for data preprocessing, analysis, and statistical testing in this study is available on GitHub: https://github.com/charlie-reynolds/Audio-Visual-RIFT-Older-Adults.

## Data Availability

The data supporting the findings of this study are publicly available through Zenodo at https://doi.org/10.5281/zenodo.23365028.

## Notes

### Competing Interest Statement

The authors have declared no competing interest.

