## Supplementary Information for "Older Adults Show Preserved Neural Signatures of Audiovisual Integration but Altered Links to Speech Comprehension"

#### Results:

*Behavioural Performance Across All Experimental Conditions.*

To characterise behavioural performance in the additional No-Tagging control condition, accuracy and response time were compared across all four conditions in older adults. Comprehension accuracy differed significantly across conditions (Fig. S1A), $F(3,117) = 3.46$, $p = .019$, $\eta_p^2 = .081$. Following Bonferroni correction, accuracy was significantly greater during Pure Tagging than No Tagging ($p = .046$), while no other pairwise comparisons reached significance. Response time also differed significantly across conditions (Fig. S1B), $F(3,117) = 39.51$, $p < .001$, $\eta_p^2 = .503$. Responses were significantly slower during No Tagging than during Task-Relevant AM, Task-Irrelevant AM, and Pure Tagging (all corrected $p < .001$).

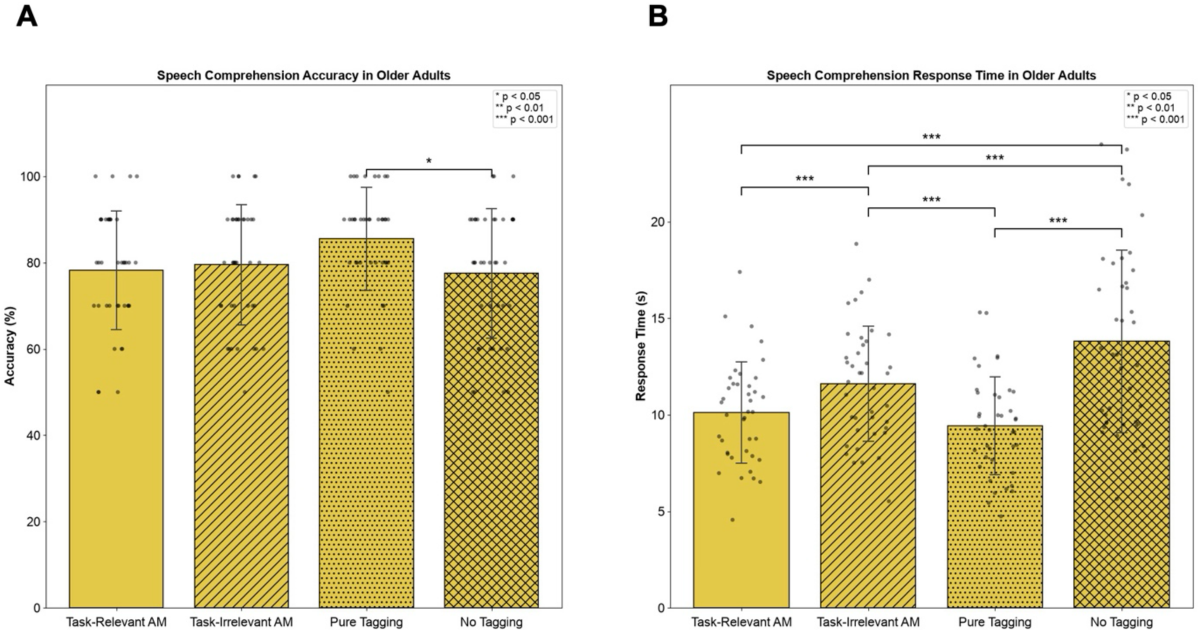


**Fig. S1 | Behavioural Performance Across All Experimental Conditions. (A)** Mean comprehension accuracy (%) in the older adult (OA) cohort across the Task-Relevant amplitude modulation (AM), Task-Irrelevant AM, Pure Tagging and No Tagging conditions. **(B)** Mean response times (RT; s) for the OA cohort across the same conditions. Error bars represent ±1 SD. Significance markers indicate Bonferroni-corrected post hoc comparisons ($p < 0.05$, $p < 0.01$, $p < 0.001$).

Group-Averaged Gaze Distributions Across Experimental Conditions

To assess whether participants maintained comparable visual attention across conditions, gaze position was examined from the first to the final Trigger 2 event in each recording. For each participant and condition, gaze samples falling within this interval were used to generate a two-dimensional gaze-density map. Individual maps were normalised before averaging across participants so that each participant contributed equally to the group-level estimate. Supplementary Figure S2 shows the resulting group-averaged gaze distributions for (A) Task-Relevant AM, (B) Task-Irrelevant AM, (C) Pure Tagging, and (D) No Tagging. Across conditions, gaze was concentrated within the central region of the display, indicating broadly consistent visual fixation throughout the task.

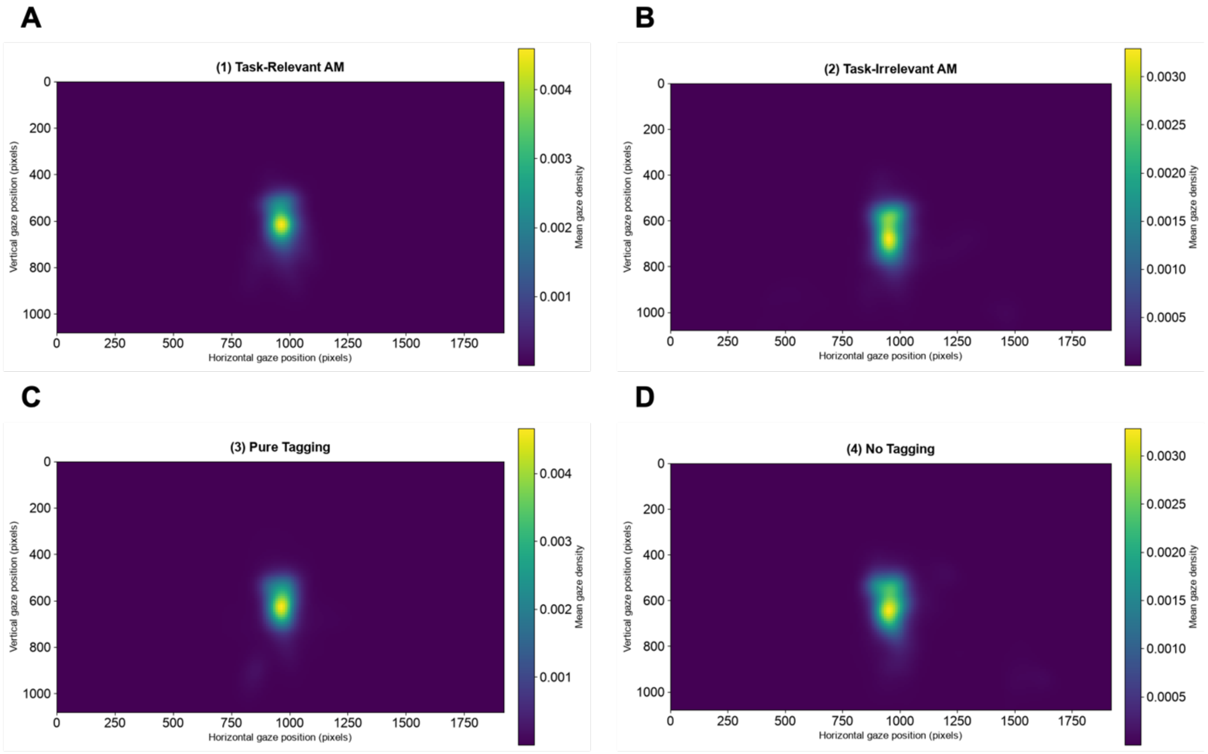


**Fig. S2 | Group-averaged gaze distributions across experimental conditions. (A)** Task-Relevant amplitude modulation (AM), **(B)** Task-Irrelevant AM, **(C)** Pure Tagging and **(D)** No Tagging. For each participant and condition, gaze samples recorded between the first and final trigger were used to generate a two-dimensional gaze-density map. Individual participant maps were normalised before averaging so that each participant contributed equally to the group-level distribution. Across all four conditions, gaze was concentrated within the central region of the display, indicating broadly comparable visual attention across conditions.

### *Complete Behavioural Statistics (Within OA Sample)*

#### Table S1. Accuracy: Bonferroni-Corrected Pairwise Comparisons

| Comparison | t(39) | Cohen's dz | p uncorrected | p Bonferroni |
|---|---|---|---|---|
| Task-Relevant AM vs Task-Irrelevant AM | -0.49 | -0.08 | 0.628 | 1.000 |
| Task-Relevant AM vs Pure Tagging | -2.44 | -0.39 | 0.019 | 0.117 |
| Task-Relevant AM vs No Tagging | 0.29 | 0.05 | 0.776 | 1.000 |
| Task-Irrelevant AM vs Pure Tagging | -2.42 | -0.38 | 0.020 | 0.121 |
| Task-Irrelevant AM vs No Tagging | 0.65 | 0.10 | 0.518 | 1.000 |
| Pure Tagging vs No Tagging | 2.81 | 0.44 | 0.008 | 0.046 |

#### Table S2. Response Time: Bonferroni-Corrected Pairwise Comparisons

| Comparison | t(39) | Cohen's dz | p uncorrected | p Bonferroni |
|---|---|---|---|---|

| | | | | |
|---|---|---|---|---|
| Task-Relevant AM vs Task-Irrelevant AM | -4.40 | -0.70 | < 0.001 | < 0.001 |
| Task-Relevant AM vs Pure Tagging | 2.17 | 0.34 | 0.036 | 0.216 |
| Task-Relevant AM vs No Tagging | -6.87 | -1.09 | < 0.001 | < 0.001 |
| Task-Irrelevant AM vs Pure Tagging | 6.27 | 0.99 | < 0.001 | < 0.001 |
| Task-Irrelevant AM vs No Tagging | -5.04 | -0.80 | < 0.001 | < 0.001 |
| Pure Tagging vs No Tagging | -7.64 | -1.21 | < 0.001 | < 0.001 |

*Complete Behavioural Statistics (Between OA and YA Sample)*

Table S3. Full mixed-design ANOVA results for comprehension accuracy.

| Source | df | F | p | Partial η² |
|---|---|---|---|---|
| Age Group | 1, 78 | 3.415 | .068 | 0.042 |
| Condition | 2, 156 | 2.051 | .132 | 0.026 |
| Age Group × Condition | 2, 156 | 8.233 | < .001 | 0.095 |

Note. Mixed ANOVA with Age Group (younger adults, older adults) as the between-subject factor and Condition (Task-Relevant AM, Task-Irrelevant AM, Pure Tagging) as the within-subject factor. $\eta^2p$ = partial eta squared.

Table S4. Full mixed-design ANOVA results for response time.

| Source | df | F | p | Partial η² |
|---|---|---|---|---|
| Age Group | 1, 78 | 17.186 | < .001 | 0.181 |
| Condition | 2, 156 | 34.972 | < .001 | 0.310 |
| Age Group × Condition | 2, 156 | 4.283 | .015 | 0.052 |

Note. Mixed ANOVA with Age Group (younger adults, older adults) as the between-subject factor and Condition (Task-Relevant AM, Task-Irrelevant AM, Pure Tagging) as the within-subject factor. $\eta^2p$ = partial eta squared.

Table S5. Between-group comparisons for comprehension accuracy.

| Condition | YA Mean ± SD (%) | OA Mean ± SD (%) | Welch t | df | Cohen's d | p uncorrected | p BH-FDR |
|---|---|---|---|---|---|---|---|
| Task-Relevant AM | 89.62 ± 9.75 | 78.25 ± 13.75 | 4.267 | 70.30 | 0.95 | < .001 | < .001 |
| Task-Irrelevant AM | 81.80 ± 14.11 | 79.50 ± 13.95 | 0.733 | 77.99 | 0.16 | .466 | .466 |
| Pure Tagging | 82.10 ± 11.43 | 85.50 ± 11.97 | -1.299 | 77.83 | -0.29 | .198 | .296 |

Note. Independent-samples Welch tests compare younger adults (YA) with older adults (OA) within each matched condition; N = 40 per group. Cohen's d is calculated as YA − OA. p values are corrected across the three condition-wise comparisons using the Benjamini–Hochberg false discovery rate (BH-FDR).

Table S6. Between-group comparisons for response time.

| Condition | YA Mean ± SD (s) | OA Mean ± SD (s) | Welch t | df | Cohen's d | p uncorrected | p BH-FDR |
|---|---|---|---|---|---|---|---|

| | | | | | | | |
|---|---|---|---|---|---|---|---|
| Task-Relevant AM | 7.59 ± 2.02 | 10.11 ± 2.63 | -4.800 | 73.25 | -1.07 | < .001 | < .001 |
| Task-Irrelevant AM | 9.35 ± 2.65 | 11.61 ± 2.98 | -3.586 | 76.96 | -0.80 | < .001 | < .001 |
| Pure Tagging | 8.18 ± 1.81 | 9.43 ± 2.52 | -2.548 | 70.83 | -0.57 | .013 | .013 |

Note. Independent-samples Welch tests compare younger adults (YA) with older adults (OA) within each matched condition; N = 40 per group. Cohen's d is calculated as YA − OA. p values are corrected across the three condition-wise comparisons using the Benjamini–Hochberg false discovery rate (BH-FDR).

*Complete MEG Cluster Statistics*

Table S7. Summary of MEG Cluster Statistics: Main Effect of Age Group

| Measure | Frequency | Direction | Significant clusters | Cluster size | Cluster-corrected p |
|---|---|---|---|---|---|
| Visual tagging | 55 Hz | — | 0 | — | No significant clusters |
| Auditory tagging | 40 Hz | OA > YA | 1 | 871 vertices | 0.018 |
| Intermodulation | 15 Hz | — | 0 | — | No significant clusters |

Statistical significance was assessed using cluster-based permutation testing. Only analyses yielding significant clusters are reported. Multiple significant clusters within a comparison are listed by their cluster-corrected p-values.

Table S8. Summary of MEG Cluster Statistics: Condition Effects in Older Adults

| Measure | Frequency | Contrast | Direction | Cluster | Cluster size (vertices) | Cluster-corrected p |
|---|---|---|---|---|---|---|
| Visual tagging | 55 Hz | TR vs TI | TR > TI | 0 | 796 | 0.002 |
| Visual tagging | 55 Hz | TR vs TI | TR > TI | 3 | 246 | 0.038 |
| Visual tagging | 55 Hz | TR vs TI | TR > TI | 5 | 334 | 0.018 |
| Visual tagging | 55 Hz | TR vs TI | TR > TI | 86 | 1808 | < 0.001 |
| Visual tagging | 55 Hz | TR vs TI | TR > TI | 87 | 792 | 0.003 |
| Visual tagging | 55 Hz | TR vs PT | TR < PT | 5 | 4076 | < 0.001 |
| Visual tagging | 55 Hz | TR vs PT | TR < PT | 7 | 492 | 0.041 |
| Visual tagging | 55 Hz | TR vs PT | TR < PT | 86 | 4525 | < 0.001 |
| Visual tagging | 55 Hz | TI vs PT | TI < PT | 0 | 6908 | < 0.001 |
| Visual tagging | 55 Hz | TI vs PT | TI < PT | 31 | 8188 | < 0.001 |
| Auditory tagging | 40 Hz | TR vs TI | TR < TI | 2 | 6845 | < 0.001 |
| Auditory tagging | 40 Hz | TR vs TI | TR < TI | 37 | 7914 | < 0.001 |
| Auditory tagging | 40 Hz | TR vs PT | TR < PT | 3 | 6898 | < 0.001 |
| Auditory tagging | 40 Hz | TR vs PT | TR < PT | 38 | 8220 | < 0.001 |
| Auditory tagging | 40 Hz | TI vs PT | — | — | — | No significant clusters |
| Intermodulation | 15 Hz | TR vs TI | TR > TI | 3 | 323 | 0.034 |
| Intermodulation | 15 Hz | TR vs TI | TR > TI | 129 | 308 | 0.035 |
| Intermodulation | 15 Hz | TR vs TI | TR > TI | 130 | 421 | 0.016 |
| Intermodulation | 15 Hz | TR vs PT | — | — | — | No significant clusters |
| Intermodulation | 15 Hz | TI vs PT | TI < PT | 141 | 648 | < 0.001 |

Note. TR = Task-Relevant AM; TI = Task-Irrelevant AM; PT = Pure Tagging. Direction is based on the sign of the cluster t-statistics for the stated contrast. Only significant clusters (cluster-corrected p < .05) are listed; contrasts with no significant clusters are retained explicitly. Cluster p values were obtained using cluster-based permutation testing.

Table S9. Summary of MEG Cluster Statistics: Condition Effects Across Age Groups

| Measure | Frequency | Contrast | Direction | Cluster | Cluster size (vertices) | Cluster-corrected p |
|---|---|---|---|---|---|---|
| Visual tagging | 55 Hz | TR vs TI | TR > TI | 1 | 477 | 0.013 |
| Visual tagging | 55 Hz | TR vs TI | TR > TI | 2 | 2548 | < 0.001 |
| Visual tagging | 55 Hz | TR vs TI | TR > TI | 107 | 2907 | < 0.001 |
| Visual tagging | 55 Hz | TR vs TI | TR > TI | 108 | 1409 | < 0.001 |
| Visual tagging | 55 Hz | TR vs PT | TR < PT | 1 | 6400 | < 0.001 |
| Visual tagging | 55 Hz | TR vs PT | TR < PT | 55 | 7023 | < 0.001 |
| Visual tagging | 55 Hz | TI vs PT | TI < PT | 1 | 9527 | < 0.001 |
| Visual tagging | 55 Hz | TI vs PT | TI < PT | 2 | 10028 | < 0.001 |
| Auditory tagging | 40 Hz | TR vs TI | TR < TI | 2 | 8445 | < 0.001 |
| Auditory tagging | 40 Hz | TR vs TI | TR < TI | 13 | 9033 | < 0.001 |
| Auditory tagging | 40 Hz | TR vs PT | TR < PT | 2 | 8456 | < 0.001 |
| Auditory tagging | 40 Hz | TR vs PT | TR < PT | 19 | 9041 | < 0.001 |
| Auditory tagging | 40 Hz | TI vs PT | — | — | — | No significant clusters |
| Intermodulation | 15 Hz | TR vs TI | TR > TI | 0 | 1595 | < 0.001 |
| Intermodulation | 15 Hz | TR vs TI | TR > TI | 1 | 2368 | < 0.001 |
| Intermodulation | 15 Hz | TR vs PT | — | — | — | No significant clusters |
| Intermodulation | 15 Hz | TI vs PT | TI < PT | 0 | 574 | 0.002 |
| Intermodulation | 15 Hz | TI vs PT | TI < PT | 1 | 1165 | < 0.001 |

Note. TR = Task-Relevant AM; TI = Task-Irrelevant AM; PT = Pure Tagging. Condition effects were assessed across the combined younger adult (YA) and older adult (OA) samples. Direction is based on the sign of the cluster statistics for the stated contrast. Only significant clusters (cluster-corrected $p < .05$) are listed; contrasts with no significant clusters are retained explicitly. Cluster p values were obtained using cluster-based permutation testing.

Table S10. Summary of MEG Cluster Statistics: Age Group × Condition Interactions

| Measure | Frequency | Contrast | Significant clusters | Minimum cluster-corrected p |
|---|---|---|---|---|
| Visual tagging | 55 Hz | TR vs TI | 0 | 0.204 |
| Visual tagging | 55 Hz | TR vs PT | 0 | 0.497 |
| Visual tagging | 55 Hz | TI vs PT | 0 | 0.350 |
| Auditory tagging | 40 Hz | TR vs TI | 0 | 0.191 |
| Auditory tagging | 40 Hz | TR vs PT | 0 | 0.032 |
| Auditory tagging | 40 Hz | TI vs PT | 0 | 0.541 |
| Intermodulation | 15 Hz | TR vs TI | 0 | 0.380 |
| Intermodulation | 15 Hz | TR vs PT | 0 | 0.127 |
| Intermodulation | 15 Hz | TI vs PT | 0 | 0.026 |

Note. TR = Task-Relevant AM; TI = Task-Irrelevant AM; PT = Pure Tagging. Age Group × Condition interactions were tested separately for each pairwise condition contrast. The significance threshold was Bonferroni-corrected across the three interaction contrasts ($\alpha = .0167$). No interaction clusters reached this corrected threshold. The minimum cluster-corrected p value is reported for each analysis.

Table S11. Summary of MEG Cluster Statistics: Behaviour-weighted intermodulation

| Analysis | Frequency | Direction | Cluster | Cluster size (vertices) | t min | t max | t mean | Cluster-corrected p |
|---|---|---|---|---|---|---|---|---|
| Behaviour-weighted Task-Relevant intermodulation | 15 Hz | YA > OA | 153 | 291 | 1.99 | 4.37 | 2.64 | 0.022 |

| | | | | | | | | |
|---|---|---|---|---|---|---|---|---|
| Behaviour-weighted Task-Relevant intermodulation | 15 Hz | YA > OA | 154 | 242 | 2.00 | 4.56 | 2.65 | 0.040 |

Note. YA = younger adults; OA = older adults. Behaviour-weighted intermodulation coherence was calculated for the Task-Relevant AM condition at 15 Hz. Positive t-values indicate greater behaviour-weighted intermodulation coherence in YA than OA. Only significant clusters (cluster-corrected $p < 0.05$) are reported.
